# Classifying Literature Extracted Events for Automated Model Extension

**DOI:** 10.1101/2021.09.30.462421

**Authors:** Casey Hansen, Julia Kisslinger, Neal Krishna, Emilee Holtzapple, Yasmine Ahmed, Natasa Miskov-Zivanov

## Abstract

In this study, we investigate the integration of three previously developed tools: FLUTE, VIOLIN, and CLARINET. We show how using these tools together adds additional capabilities in extending models from relevant research literature. We illustrate how we plan to address current modeling pitfalls with these tools (such as machine reading errors and literature volume), and how we plan to use these tools as the foundation for an automated model extension framework. Documentation and links to our tools can be found at:

violin-tool.readthedocs.io
clarinet-docs.readthedocs.io
flute.readthedocs.io

## 2 INTRODUCTION

Machine reading tools are able to quickly and automatically curate vast amounts of information from relevant published literature [6][2]. This curated information can be used to build biological computational models or expand upon existing models. However, the information gleaned by machine readers is both vast and varied in quality. Machine readers must work to extract standardized biological interactions from inconsistent terminology and complex sentence structures, which sometimes leads to extraction errors.

Previously we have developed VIOLIN (Verifying Interactions of Likely Importance to the Network) a tool to automatically classify and judge biological interactions extracted from relevant literature. With VIOLIN, we are able to take these literature extracted events (LEEs) and compare them to an existing biological model, determining whether a given LEE agrees with the model (corroborates), introduces new information to the model (extends), disputes the model (contradicts), or requires manual review (flagged). Each LEE is assigned four numerical values to represent its relationship to the model system (Match Score), its classification category (Kind Score), its frequency (Evidence Score), and extraction confidence (Epistemic Value). These values are combined into a Total Score to allow for automatic filtering and classification of large sets of LEEs curated from multiple sources. To further increase the utility of VIOLIN, we now seek to integrate VIOLIN as part of an automated model-building framework (Figure 1).

**Figure 1:**
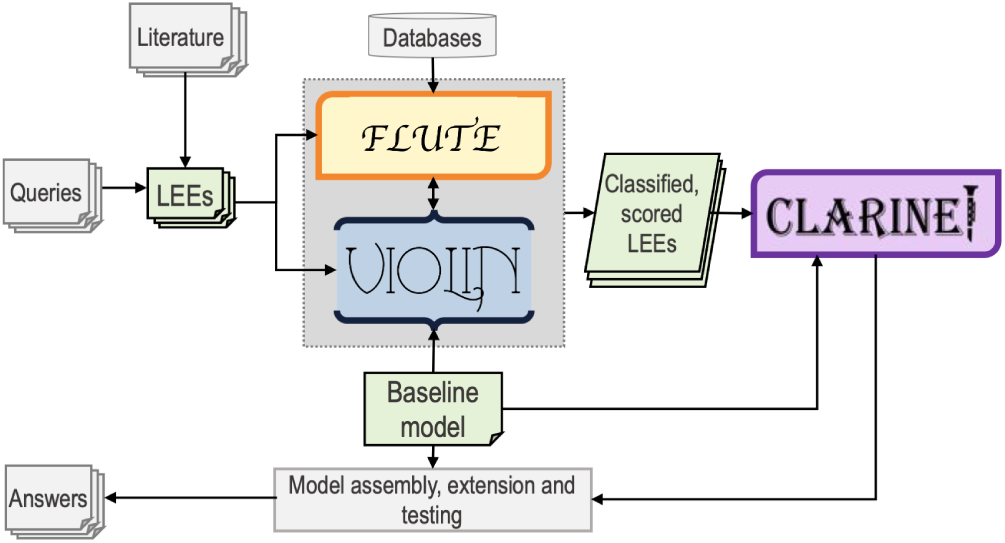
An outline of automated information extraction and the model assembly, highlighting the roles FLUTE, VIOLIN, and CLARINET hold: FLUTE and VIOLIN judge the quality, relevance, and usefulness of LEEs on an individual basis, and then CLARINET judges how the LEEs connect, both to each other and the baseline model.

Current approaches towards building and extending models have two major pitfalls. They either focus on only a single step of the process [8][7], or the decision metrics lack depth, focusing on the machine reading output or model as separate entities, more than the relationship between the two [2].

We first integrated VIOLIN with the filtering tool FLUTE (FiLter for Understanding True Events) [3], to make use of the expert data gathered in public databases. The FLUTE tool connects to public protein interaction databases to judge the accuracy of an LEE. This integration allows us to balance the removal of erroneous extractions while retaining novel interactions which may not yet be represented in a database.

We next integrated VIOLIN with CLARINET (CLARIfying NET-works) [1], an automated model extension tool. Where VIOLIN classifies individual LEEs for their relevance and usefulness to a given model, CLARINET classifies candidate extensions as clusters of biological interactions, taking into account how LEEs are connected to each other, in addition to their connection to the baseline model.

These three tools together create a powerful method of taking information-rich relevant literature and identifying the highest quality events for extending a baseline model.

## 3 METHODS

To evaluate the itnegration of VIOLIN and FLUTE, we used the following as inputs (1) three computational models, namely a model of Skel-133 Melanoma, a model of human T-cells [4], and a model of the BDNF pathway as it relates to Major Depressive Disorder (MDD) [5], and (2) four LEE sets for each model. From these inputs, we generated three types of outputs: LEE sets classified by VIOLIN only (control), LEE sets first filtered by FLUTE and then classified by VIOLIN (pre-processed), and LEE sets first classified by VIOLIN and then filtered by FLUTE (post-processed).

We next investigated the integration of VIOLIN and CLARINET, using a Glioblastoma Multiforme model and two highly specialized LEE sets. Our first LEE set (R_*G*1_), contained 10,130 LEEs from 242 papers, and the second (R_*G*2_) contained 25,875 LEEs from 454 papers. From these inputs, we also created three data outputs: candidate clusters created from the raw LEE sets (control), candidate clusters created from the Total VIOLIN output, which lists only the unique LEEs (unique), and candidate clusters created from only the VIOLIN extensions (extensions). Table 1 shows a summary of the input parameters for both parts of our investigation.

**Table 1:** Testing Inputs.

| LEE Suffix | Model | Model Nodes | LEE sets |
| --- | --- | --- | --- |
| $R_A$ | T cell | 61 | 4 |
| $R_B$ | Melanoma | 225 | 4 |
| $R_C$ | MDD | 72 | 4 |
| $R_G$ | GBM | 238 | 2 |

## 4 RESULTS

For our VIOLIN-FLUTE integration, we found that post-processing methods had interesting implications for the VIOLIN classification categories (Figure 2). Post-processing methods work as a feed-back for VIOLIN, showing how machine reading errors propagate through VIOLIN’s judgement, and also helping drive the user’s choices for choosing LEEs. Contradictions and Extensions had the lowest average retention rate, and flagged had the highest (Table 2). This supports our previous suggestion that the contradiction category can be used to filter out machine reading errors. As expected, those LEEs with high evidence scores are retained more often than those with low evidence scores.

**Figure 2:**
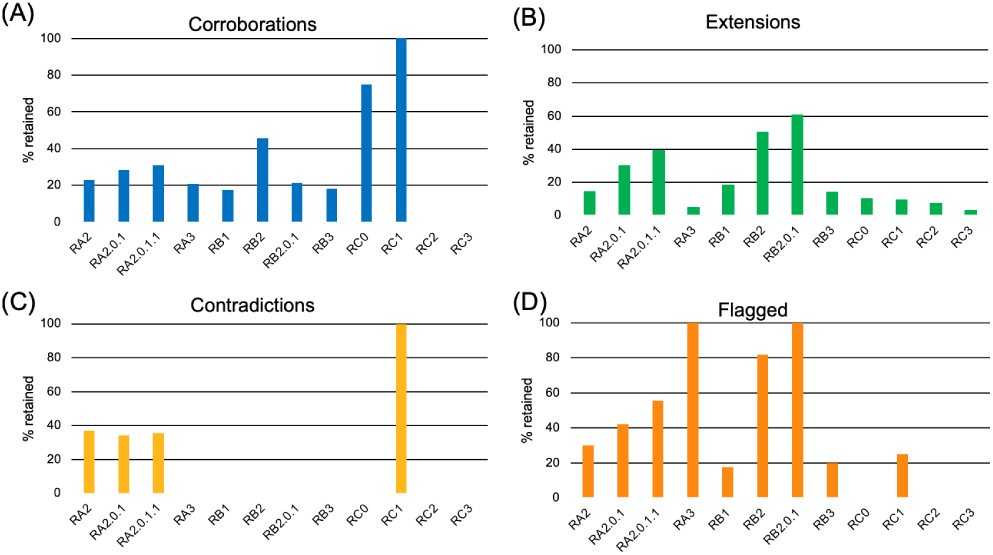
Retention counts for each VIOLIN classification: (A) shows corroborations, (B) shows extensions, (C) shows contradictions, and (D) shows flagged

**Table 2:** Average Retention Rates.

|  | Average % Retained |
| --- | --- |
| Corroborations | 32.9 |
| Extensions | 21.7 |
| Contradictions | 10.7 |
| Flagged | 39.3 |

For the VIOLIN-CLARINET integration, we observed the size and central nodes of the candidate clusters for the control, unique, and extension output from CLARINET (Figure 3). We found that just the act of comparing the control output to the unique output, which contains only single instances of a given LEE, has an effect on the outcome of the candidate clusters. The candidate clusters from the extensions are an even more focused input, as they only present LEEs for consideration which are known to present new information to the model. This suggests that forming candidate clusters from raw machine reading output is influenced by corroborative or contradictory LEEs, as well as machine reading errors, and having more directed LEE sets would produce more directed clusters.

**Figure 3:**
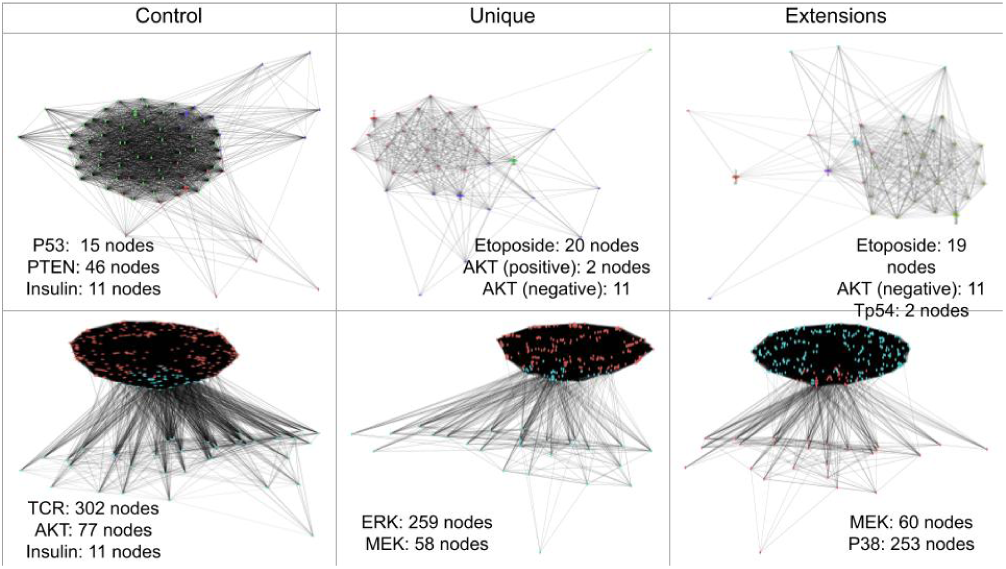
Candidate clusters for the control, unique, and extensions input compared to the GBM model using CLARINET. The top row was created from the R_*G*1_ LEE set, and the bottom row was created from the R_*G*2_ LEE set

## 5 CONCLUSIONS

Integrating VIOLIN with FLUTE and CLARINET showed us the promising outcome of combining these individually effective tools. The options of using FLUTE with VIOLIN allows the user to determine the importance of removing erroneous information versus retaining novel interactions. Our results from CLARINET show that narrowing an LEE set down to those which are most useful for extension changes the candidate extensions, and VIOLIN allows this process to be fast and automatic. Our next step is to further investigate approaches to utilize the integration of these three tools towards automated creation of useful and reliable models.

## ACKNLOWEDGEMENTS

This project was funded by DARPA award W911NF-17-1-0135.

## REFERENCES

[1] Ahmed, Y., Telmer, C., and Miskov-Zivanov, N. CLARINET: Efficient learning of dynamic network models from literature.

[2] Gyori, B. M., Bachman, J. A., Subramanian, K., Muhlich, J. L., Galescu, L., and Sorger, P. K. From word models to executable models of signaling networks using automated assembly. 954.

[3] Holtzapple, E., Telmer, C. A., and Miskov-Zivanov, N. FLUTE: Fast and reliable knowledge retrieval from biomedi-cal literature. _eprint: https://academic.oup.com/database/article-pdf/doi/10.1093/database/baaa056/33571972/baaa056.pdf.

[4] Miskov-Zivanov, N., Turner, M. S., Kane, L. P., Morel, P. A., and Faeder, J. R. The duration of t cell stimulation is a critical determinant of cell fate and plasticity. ra97–ra97.

[5] Sandhya, V. K., Raju, R., Verma, R., Advani, J., Sharma, R., Radhakrishnan, A., Nanjappa, V., Narayana, J., Somani, B. L., Mukherjee, K. K., Pandey, A., Christopher, R., and Prasad, T. S. K. A network map of BDNF/TRKB and BDNF/p75ntr signaling system. 301–307.

[6] Valenzuela-Escárcega, M. A., Hahn-Powell, G., Surdeanu, M., and Hicks, T. A domain-independent rule-based framework for event extraction. In Proceedings of ACL-IJCNLP 2015 System Demonstrations (Beijing, China, July 2015), Association for Computational Linguistics and The Asian Federation of Natural Language Processing, pp. 127–132.

[7] von Mering, C., Jensen, L. J., Snel, B., Hooper, S. D., Krupp, M., Foglierini, M., Jouffre, N., Huynen, M. A., and Bork, P. STRING: known and predicted protein-protein associations, integrated and transferred across organisms. Nucleic Acids Res 33, Database issue (Jan 2005), D433–437.

[8] Zerva, C., Batista-Navarro, R., Day, P., and Ananiadou, S. Using uncertainty to link and rank evidence from biomedical literature for model curation. Bioinformatics 33, 23 (Dec 2017), 3784–3792.

